# First large-scale peach gene coexpression network: A new tool for predicting gene function

**DOI:** 10.1101/2023.06.22.546058

**Authors:** Felipe Pérez de los Cobos, Beatriz E. García-Gómez, Luis Orduña-Rubio, Ignasi Batlle, Pere Arús, José Tomás Matus, Iban Eduardo

**Affiliations:** Institut de Recerca i Tecnologia Agroalimentàries (IRTA), Mas Bové, Ctra. Reus-El Morell Km 3,8 43120 Constantí Tarragona, Spain; Institut de Recerca i Tecnologia Agroalimentàries (IRTA), Centre de Recerca en Agrigenòmica (CRAG), CSIC-IRTA-UAB-UB. Cerdanyola del Vallès (Bellaterra), 08193 Barcelona, Spain; Centre for Research in Agricultural Genomics (CRAG) CSIC-IRTA-UAB-UB, Campus UAB, Edifici CRAG, Cerdanyola del Vallès (Bellaterra), 08193 Barcelona, Spain; Institute for Integrative Systems Biology (I2SysBio), Universitat de Valencia-CSIC, Paterna, 46908, Valencia, Spain

## Abstract

Transcriptomics studies generate enormous amounts of biological information. Nowadays, representing this complex data as gene coexpression networks (GCNs) is becoming commonplace. Peach is a model for Prunus genetics and genomics, but identifying and validating genes associated to peach breeding traits is a complex task. A GCN capable of capturing stable gene-gene relationships would help researchers overcome the intrinsic limitations of peach genetics and genomics approaches and outline future research opportunities. In this study, we created the first large-scale GCN in peach, applying aggregated and non-aggregated methods to create four GCNs from 604 Illumina RNA-Seq libraries. We evaluated the performance of every GCN in predicting functional annotations using a machine-learning algorithm based on the ‘guilty-by-association’ principle. The GCN with the best performance was COO300, encompassing 21,956 genes and an average AUROC of 0.746. To validate its performance predicting gene function, we used two well-characterized genes involved in fruit flesh softening in peach: the endopolygalacturonases *PpPG21* and *PpPG22*. Genes coexpressing with *PpPG21* and *PpPG22* were extracted and named as melting flesh (MF) subnetwork. Finally, we performed an enrichment analysis of MF subnetwork and compared the results with the current knowledge regarding peach fruit softening process. The MF subnetwork mainly included genes involved in cell wall expansion and remodeling, with expression triggered by ripening-related phytohormones such as ethylene, auxin and methyl jasmonates. All these processes are closely related with peach fruit softening and therefore related to the function of *PpPG21* and *PpPG22*. These results validate COO300 as a powerful tool for peach and *Prunus* research. COO300, renamed as PeachGCN v1.0, and the scripts necessary to perform a function prediction analysis using it, are available at https://github.com/felipecobos/PeachGCN.

## INTRODUCTION

The advent of omics technologies has allowed the scientific community to generate enormous amounts of biological information. In parallel, increasingly efficient bioinformatic tools help us transform this information into structured biological knowledge. To date, more than seven million RNA-Seq libraries are available at the National Center of Biotechnology Information (NCBI, https://www.ncbi.nlm.nih.gov/), representing a great opportunity for large-scale bioinformatics analysis and biological data integration. Therefore, taking advantage of this valuable resource is essential in the age of big data analytics.

In transcriptomics, representing this complex data as gene coexpression networks (GCNs) is becoming a widespread practice. GCNs are usually represented as undirected graphs, where nodes correspond to genes and edges correspond to correlations in expression patterns of genes. GCNs can be built across multiple experimental conditions (condition-independent GCNs) or in specific experimental conditions (condition-dependent GCNs, e.g., tissue specific GCNs). They are based on the ‘guilt-by-association’ (GBA) principle (Oliver, 2000), which states that genes with related functions share similar expression patterns. Following this principle, and using the functional annotation of the genes forming the network, GCNs can be a very powerful tool to infer potential gene functions to specific genes or gene families and to understand the regulation of specific metabolic pathways. For this reason, GCNs are extremely useful in crop species, where most of the bioinformatic and genetic tools are modest and our understanding of gene function is still limited (Schaefer et al., 2017). Several studies have already created GCNs in the plant model *Arabidopsis thaliana* (Amrine, Blanco-Ulate, & Cantu, 2015; Furuya et al., 2021; Liu et al., 2019; Mao, Van Hemert, Dash, & Dickerson, 2009), maize (Huang et al., 2017; Ma et al., 2017), rice (Childs et al., 2011; Ficklin et al., 2010), wheat (Lv et al., 2020) and grapevine (Orduña et al., 2022; Orduña-Rubio et al., 2023; Wong, 2020; Wong et al., 2016).

Peach [*Prunus persica* L. (Batsch)] has been used as a model organism for genetics and genomics in the *Rosaceae*, and more specifically in the *Prunus* genus, which also encompasses other crops such as sweet and tart cherry, European and Japanese plum, apricot and almond. However, in peach, the validation of genes responsible for breeding traits is a complex task. Long intergeneration times and phenological cycles and space constraints due to the large size of the individuals under study are some of the hindrances for the work of peach geneticists (Aranzana et al., 2019). Moreover, there is a lack of efficient genetic transformation systems (Limera et al., 2017; Ricci et al., 2020). As a result of these limitations, only two genes to date, DRO1 and TAC1, have been biologically validated based on mutant analysis (Dardick et al., 2013; Guseman et al., 2017).

Although small-scale condition-dependent GCNs have been reported in peach and other *Prunus* species (García-Gómez et al., 2020; Jiang et al., 2023; Wang et al., 2023; Wu et al., 2021; Xi, Feng, Liu, Zhang, & Zhao, 2019; Zhang et al., 2019), these were created ad-hoc to study specific biological processes and so cannot be used in other experimental contexts. Therefore, a GCN capable of capturing robust gene-gene relationships under different experimental conditions, developmental stages and tissues is needed. A GCN with these characteristics will help researchers overcome the intrinsic limitations of peach genetics and genomics approaches and outline future research opportunities.

In this study, we present the first large-scale GCN in peach. We constructed four GCNs from publicly available RNA-Seq data and evaluated the performance of every GCN using a machine-learning algorithm based on the GBA principle. The GCN with the best performance was validated by predicting gene functions of well-characterized genes. Finally, we provide the scripts and data needed for function prediction analyses using the GCN presented in this study. These resources can be found at https://github.com/felipecobos/PeachGCN.

## MATERIALS AND METHODS

### Data compilation

Forty-nine independent Sequence Read Archive (SRA) Bioprojects, encompassing 608 RNA-Seq libraries (Supplementary Material 1) were downloaded from the SRA database (Leinonen et al., 2011) in the NCBI (Sayers et al., 2022a). These RNA-Seq libraries represented all the libraries available in the NCBI to date 09/04/2020. The peach reference genome ‘Lovell’ version 2.1 (Verde et al., 2013, 2017) and its functional annotation were downloaded from Genome Database for *Rosaceae* (GDR) (Jung et al., 2019). Finally, seven functional gene annotation datasets were retrieved using the methods described below. Gene Ontology peach functional terms for biological process (GObp), molecular function (GOmf) and cellular component (GOcc) (Ashburner et al., 2000; Carbon et al., 2021)) and Pfam database peach classification (Mistry et al., 2021) were retrieved using the biomaRt R package (Durinck et al., 2009). Kyoto Encyclopedia of Genes and Genomes (KEGG) peach pathway annotations (Kanehisa & Goto, 2000) were retrieved using the KEGG API (https://www.kegg.jp/kegg/rest/keggapi.html). PANTHER HMM peach classifications version 16 (Mi et al., 2021) and MapMan Pathways version 4.2 (Thimm et al., 2004) were downloaded from the public repositories.

### Mapping and quality filtering

We performed a sequencing-quality filtering and adapter removal using Trim Galore! version 0.6.1 (https://www.bioinformatics.babraham.ac.uk/projects/trim_galore/). Reads with terminal Ns were trimmed, then reads with a Phred score lower than 28 or smaller than 35 nucleotides were filtered. Filtered libraries were quality checked using FastQC version 0.11.5 (http://www.bioinformatics.babraham.ac.uk/projects/fastqc/). HISAT2 version 2.1 (Kim et al., 2015) was used to map RNA sequencing libraries to the reference peach genome ‘Lovell’ version 2.1 (Verde et al., 2013, 2017) with default parameters. Mapped Binary Alignment Map (BAM) files were filtered by alignment quality using SAMtools version 1.9 (Danecek et al., 2021; Li et al., 2009). Reads with mapping quality lower than 40 were filtered out. After this filtering, BAM files with less than 5,000,000 reads were discarded, leaving a total of 498 RNA-Seq libraries from 43 independent Bioprojects for further analyses.

### Aggregated and non-aggregated GCNs inference

A raw count matrix was calculated using featureCounts (Liao et al., 2014), from Subread R package version 2.0.0 (http://subread.sourceforge.net/). For the raw count matrix construction, we excluded chimeric fragments and we used the coding DNA sequences as feature type and gene IDs as attribute type. The raw count matrix was then normalized to fragments per Kilobase million (FPKM) mapped fragments (Z. Wang, Gerstein, & Snyder, 2009), obtaining a FPKM matrix. We then applied two different methodologies: aggregated and non-aggregated network inference with two sparsity thresholds set at top 100 (stringent threshold) and 300 (relaxed threshold) ranked genes (Supplementary Figure 1).

For non-aggregated analysis, genes with less than 0.5 FPKM in 50% of the RNA-Seq libraries were removed. Pearson’s correlation coefficient (PCC) was calculated for the remaining genes and ranked according to descending PCC, giving a PCC matrix. High reciprocal rank networks for the top 100 (HRR100) and top 300 (HRR300) were constructed according to the formula:

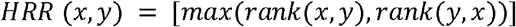

Whereby *rank(x, y)* is the descending sorted rank of gene *y* according to the coexpression list of gene *x* and vice versa for *rank(y, x)*.

For aggregated analysis, we clustered the samples into 43 different groups according to the Bioproject study ID. We filtered Bioprojects with less than six RNA-Seq libraries, leaving 26 different groups with a total of 450 RNA-Seq libraries. Genes with less than 0.5 FPKM in 50% of the libraries within each group were removed and from each filtered FPKM matrix, a high reciprocal rank network for the top 100 and top 300 was constructed. Frequency of gene coexpression interactions in all groups was calculated and ranked in a co-occurrence matrix. Finally, co-occurrence networks for top 100 (COO100) and top 300 (COO300) interactions were obtained.

### Networks performance assay

Networks were evaluated for their ability to connect peach genes sharing functional annotations. For this purpose, GBA neighbor voting, a machine learning algorithm based on the GBA principle (Ballouz et al., 2017), was assessed over the GObp, GOmf, GOcc, Pfam, KEGG, PANTHER and MapMan datasets. Each network was scored by the area under the receiver operator characteristic curve (AUROC) across all functional categories annotated for the seven datasets. Annotations were limited to groups containing 20-1,000 genes to ensure robustness and stable performance when using neighbor voting. The AUROC value threshold for an acceptable network functional annotation was set at 0.7.

We also evaluated the impact of adding individual Bioprojects to the different networks created, HRR300, HRR100, COO300 and COO100. For this purpose, we selected five subsets each of two Bioprojects computing the top 100 and top 300 HRR and COO GCNs, evaluating their AUROC using GObp, GOmf, GOcc and MapMan datasets. We repeated this process adding one Bioproject to the initial subset to reach five subsets each with 26 Bioprojects, the maximum number of Bioprojects used in this study. The final subsets corresponded to the full HRR300, HRR100, COO300 and COO100.

### Network validation

To validate the performance of COO300 in predicting gene functional annotations, we selected two well-characterized genes responsible for fruit flesh softening in peach, the endopolygalacturonases *PpPG21* and *PpPG22*, located on chromosome 4 (Table 1) (Cheng et al., 2022; Gu et al., 2016; Jiang et al., 2020; Nakano et al., 2020; Qian et al., 2021; Zhu et al., 2017). Based on the evidence available to date, the variability of flesh softening and stone adhesion during fruit ripening is due to the allelic combination of these two homologous genes. Both genes, *PpPG21* or *PpPG22*, are associated with the development of melting, non-melting or non-softening fruits, while *PpPG22* is associated with the development of freestone or clingstone fruits.

**Table 1.**
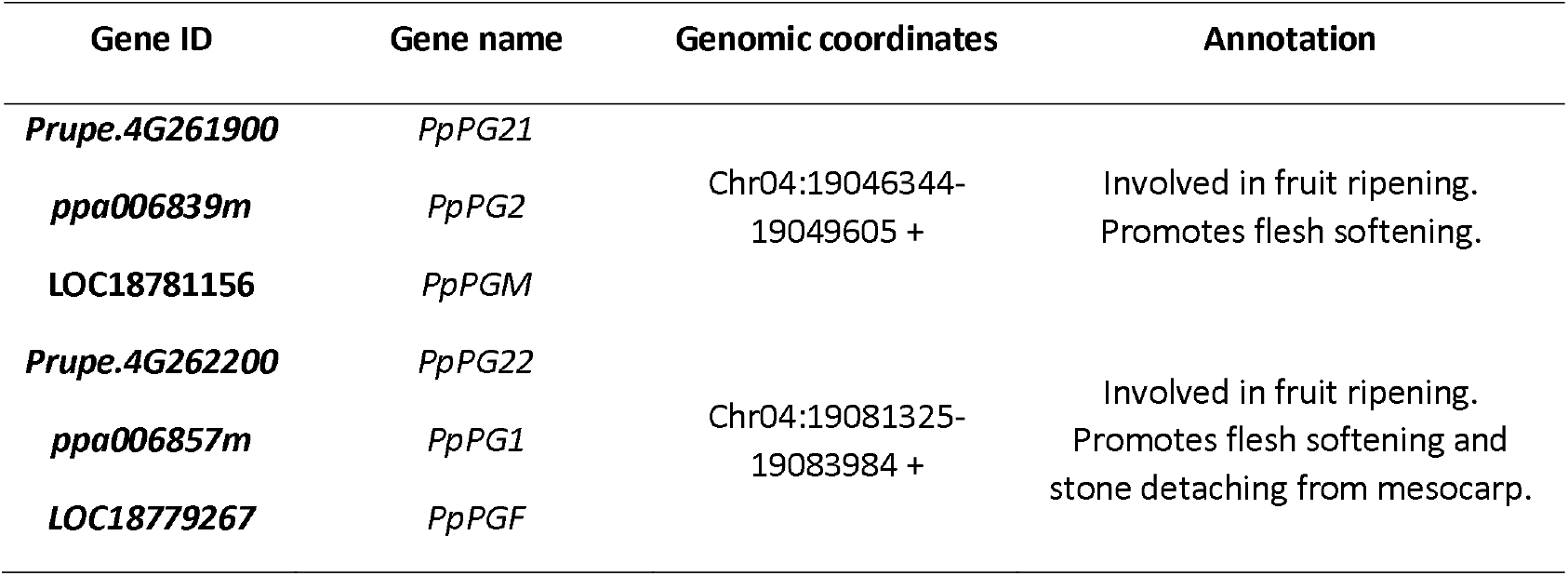
Candidate genes selected for network validation. The gene IDs were referred to the peach reference genome version 1 and 2.0 (Verde et al., 2013, 2017) and NCBI (Sayers et al., 2022b) while genomic coordinates and annotation were referred to the peach reference genome version 2.0 (Verde et al., 2017).

Genes coexpressed with *PpPG21* and *PpPG22* were extracted. Since both genes are involved in the peach fruit flesh softening process, we selected genes present in both subnetworks. The selected subnetwork, named melting flesh (MF) subnetwork, had 238 genes. With an enrichment analysis of the MF subnetwork using GObp, GOmf, GOcc and Mapman datasets we were able to identify the functional annotations statistically over-represented in each of the subnetworks studied. The significance threshold was held at *q*-value < 0.05. Finally, we compared the enriched terms (the functional annotations statistically over-represented) of the MF subnetwork with the current knowledge on the peach fruit softening process. In addition, as a negative control, we created 20 subnetworks with 238 randomly selected genes from COO300 and carried out an enrichment analysis of all negative control subnetworks following the steps described above.

## RESULTS

### Aggregated GCNs had 21,956 genes, 81.7% of the protein-coding genes annotated in the peach reference genome

To understand the differences between the GCNs, we analyzed the general topological characteristics of the four GCNs inferred in this study (Table 2). The two aggregated GCNs (COO100 and COO300) had 21,956 genes, while the two built by non-aggregated methods (HRR100 and HRR300) had 17,505 genes. Of the total number of 26,873 protein-coding genes annotated in the peach reference genome, this represented 81.7 % for aggregated and 65.1 % for non-aggregated networks. The number of genes conforming the aggregated GCNs represented 16.6 % more genes (4,451) from the peach whole-genome annotation than non-aggregated GCNs.

**Table 2.** General topological characteristics of non-aggregated and aggregated GCNs with 100 and 300 top coexpressed genes (HRR100, HRR300, COO100 and COO300).

| GCN | Number of genes | *P. persica* genes included in the GCN (%) | Range of node degree connectivity (min-max) | Average node degree connectivity |
| --- | --- | --- | --- | --- |
| **HRR100** | 17,505 | 65.1 | 649 (100-749) | 161 |
| **HRR300** | 17,505 | 65.1 | 1490 (300-1790) | 470 |
| **COO100** | 21,956 | 81.7 | 315 (100-415) | 149 |
| **COO300** | 21,956 | 81.7 | 785 (300-1085) | 442 |

The different methods used not only affected the number of genes included in the network, but also the node degree connectivity (number of coexpressed genes by gene) across all nodes of the GCN. Average node degree connectivity was higher in networks with relaxed sparsity (442 in COO300 and 470 in HRR300) in comparison to stringent sparsity (149 in COO100 and 161 in HRR100). The range between minimum and maximum node degree connectivity is wider in non-aggregated GCNs compared to aggregated GCNs with the same sparsity threshold (comparing HRR300 with COO300 and HRR100 with COO100). The minimum node degree connectivity was set by the sparsity threshold in all the networks: 100 for stringent sparsity (HRR100 and COO100) and 300 for relaxed sparsity (HRR300 and COO300). The highest node degree connectivity was found in HRR300, with a maximum of 1790 coexpressed genes with one single gene. In addition, aggregated GCNs showed a bimodal node degree connectivity distribution while non-aggregated GCNs had a unimodal distribution (Figure 1).

**Figure 1.**
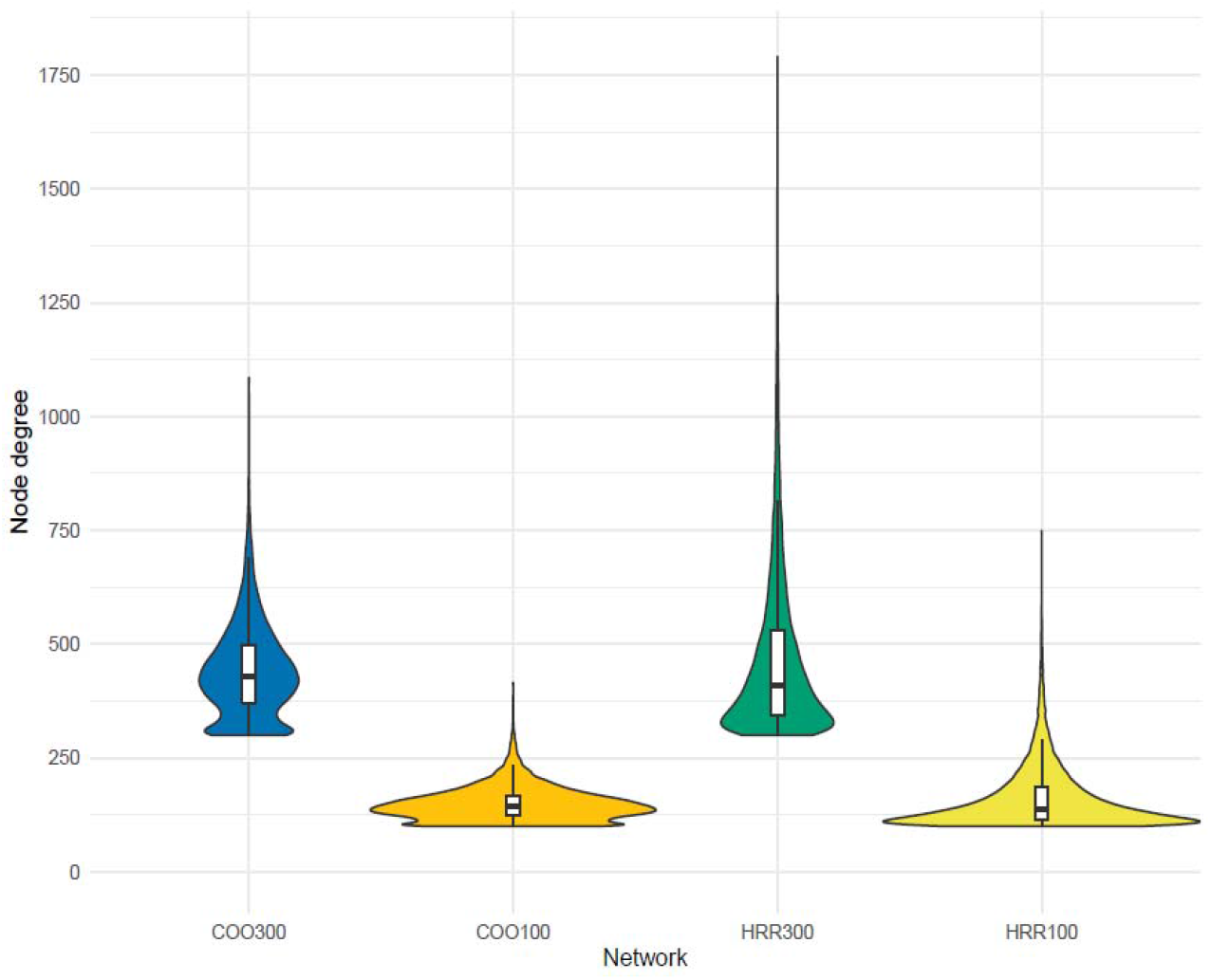
Violin plot of node degree connectivity in each of the aggregated and non-aggregated networks with relaxed or stringent sparsity (COO300, COO100, HRR300 and HRR100). Boxplots of node degree connectivity were added for each violin plot.

### COO300 was the GCN with the highest AUROC value

When considering sparsity threshold, both GCNs with relaxed sparsity (HRR300 and COO300) had AUROC values over 0.7 for all the databases annotated (Table 3). COO300 was the GCN with the highest average AUROC value (0.746), outperforming the other GCNs. COO300 had the highest mean AUROC in almost all the datasets, except for Pfam and PANTHER, where the performance of HRR300 was better than that of COO300. HRR100 and COO100 had AUROC values under 0.7 in almost all the functional annotation databases, except for the GOcc and PANTHER datasets. The best functional annotation performance in all the networks was for the functional annotation GOcc with an average AUROC value of 0.761, followed by PANTHER (0.724), KEGG (0.718), Pfam (0.714), GObp (0.709), Mapman (0.706) and GOmf (0.693).

**Table 3.** AUROC values for each GCN (COO300, HRR300, COO100, HRR100) performance in the different datasets. The best performance by dataset was highlighted with an asterisk.

| GCN | GObp | GOMf | GOcc | Pfam | KEGG | PANTHER | MapMan | Average |
| --- | --- | --- | --- | --- | --- | --- | --- | --- |
| COO300 | 0.738* | 0.723* | 0.788* | 0.736 | 0.750* | 0.746 | 0.741* | 0.746* |
| HRR300 | 0.724 | 0.705 | 0.773 | 0.745* | 0.728 | 0.749* | 0.732 | 0.736 |
| COO100 | 0.681 | 0.670 | 0.733 | 0.680 | 0.697 | 0.688 | 0.664 | 0.687 |
| HRR100 | 0.692 | 0.673 | 0.748 | 0.695 | 0.695 | 0.712 | 0.686 | 0.700 |
| Average | 0.709 | 0.693 | 0.761 | 0.714 | 0.718 | 0.724 | 0.706 | 0.717 |

The method used for network building also affected its performance, but the effect was not consistent. Considering the effect of the sparsity threshold, average AUROC values for relaxed sparsity threshold were always higher (lllHRR300 and COO300 = 0.741) than for the stringent threshold (HRR100 and COO100 = 0.694). When comparing GCNs by aggregation method, at relaxed sparsity (HRR300 and COO300), the average AUROC value for the aggregated method was higher but comparing at the stringent threshold (HRR100 and COO100), the average AUROC value was better for the non-aggregated method.

Finally, we evaluated the effects of adding Bioprojects on the AUROC value in every GCN built in this study. Figure 2 shows the correlation between the network AUROC value and the number of Bioprojects used. For every combination of GCN building method (aggregated or non-aggregated), threshold (top 300 or top 100) and dataset used (GObp, GOmf, GOcc and MapMan) we observed similar trends, where the AUROC value increased with the number of Bioprojects. This trend was more pronounced for aggregated GCNs than non-aggregated GCNs, reaching a plateau after adding 10-12 Bioprojects. In all cases, the standard deviation of aggregated GCNs decreased as the number of Bioprojects increased.

**Figure 2.**
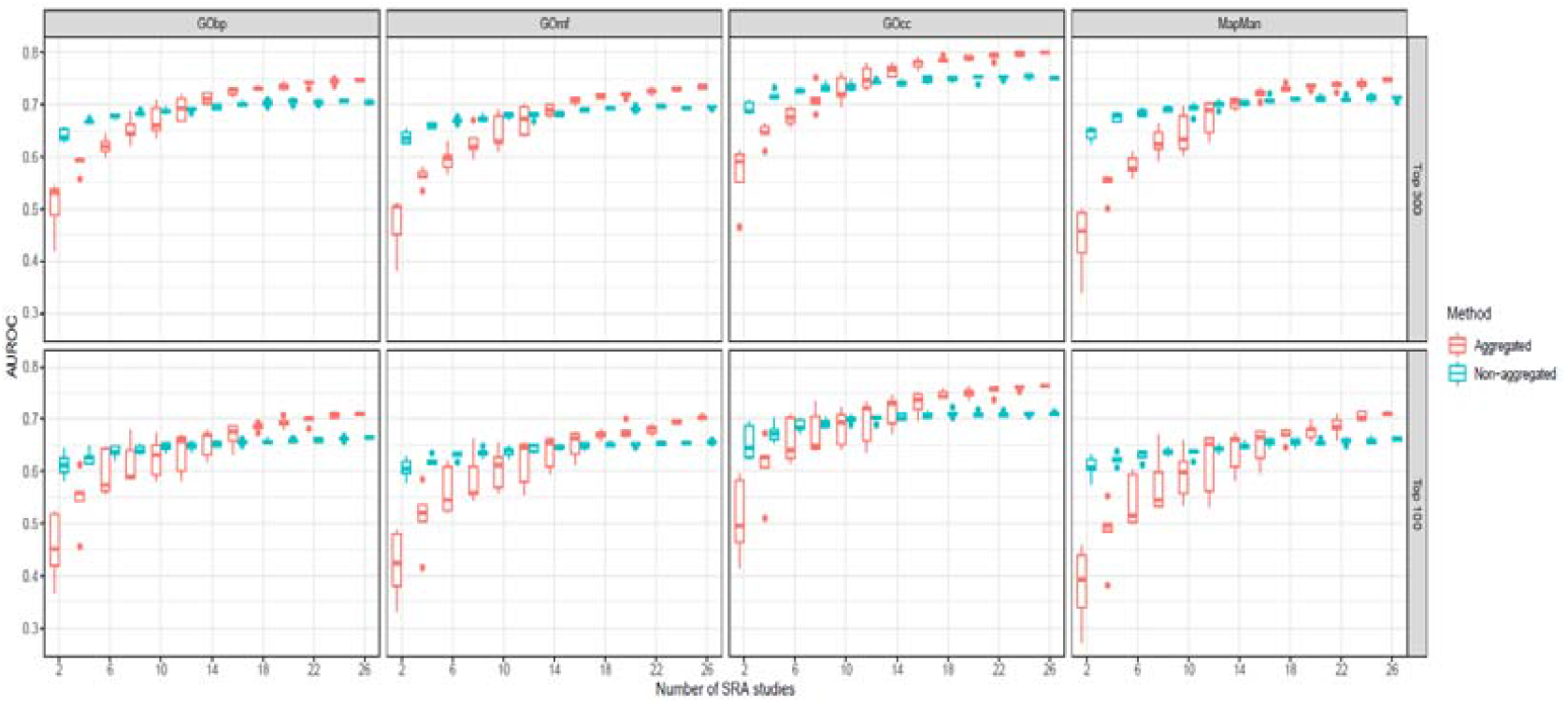
Boxplots of the AUROC value for every subset of Bioprojects (from 2 to 26) and method used.

### Aggregated GCNs showed a positive trend between average node degree connectivity and AUROC score of individual functional annotations for GObp, GOmf, GOcc, KEGG and Mapman

To assess the relationship between the AUROC score of individual functional annotations and the average node degree connectivity of the genes sharing that annotation we used a Loess regression (Figure 3). For example, an individual functional annotation could be *GOcc: cell wall*, we studied if the individual AUROC score of *GOcc: cell wall* was related to the average number of connections of the genes sharing *GOcc: cell wall* annotation. We then repeated the analysis for all the functional annotations within a dataset (*GOcc: apoplast, GOcc: extracellular region*, etc). In the case of aggregated GCNs, there was a positive trend between average node degree connectivity and AUROC score of individual functional annotations for GObp, GOmf, GOcc, KEGG and Mapman. In the case of non-aggregated GCNs the only dataset with a positive trend between average node degree connectivity and AUROC score of individual functional annotations was KEGG. The average node degree connectivity had no effect on the AUROC score of individual functional annotations in the Pfam dataset in any of the GCNs studied.

**Figure 3.**
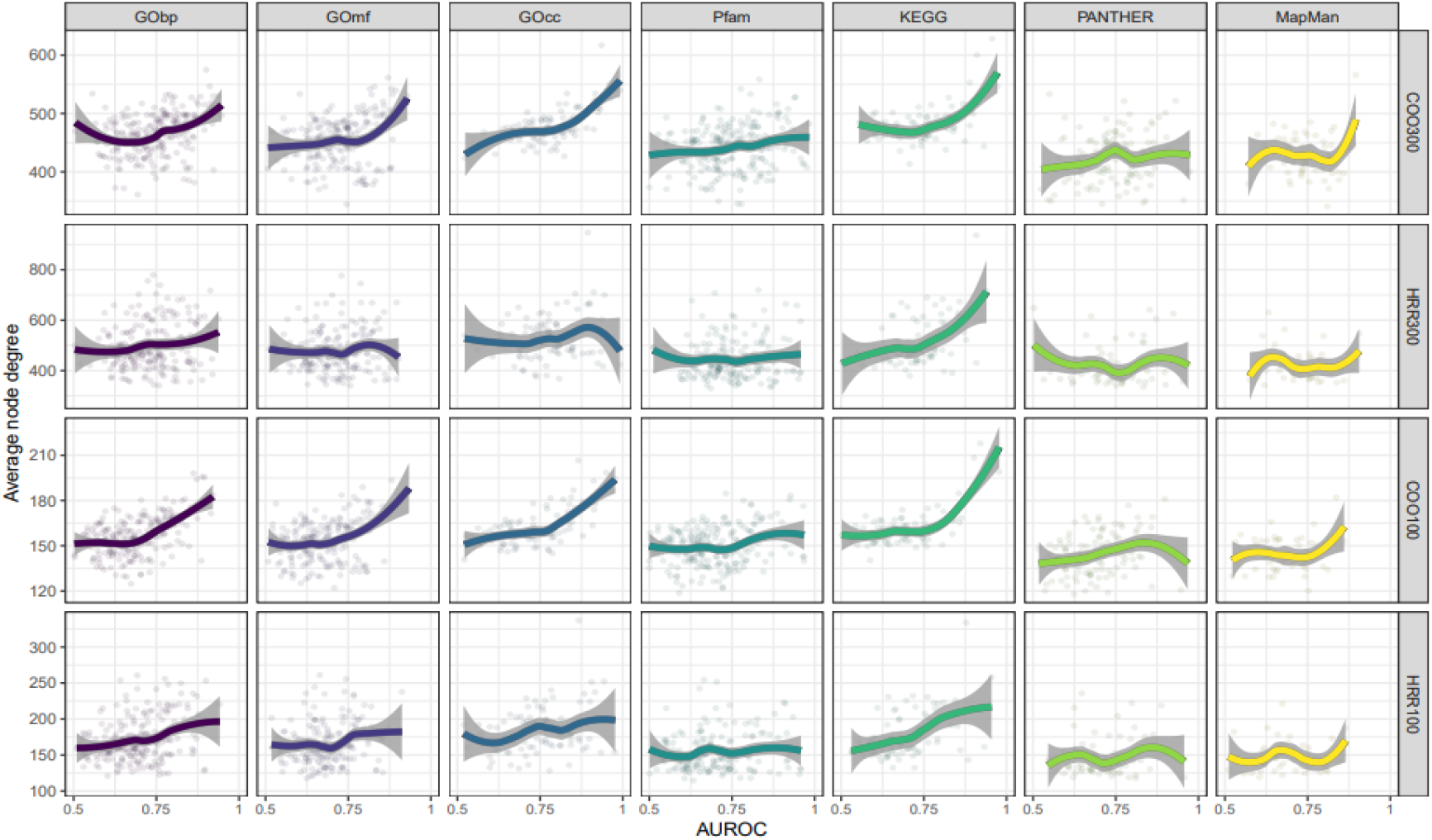
Scatter plot and Loess regression representation of average node degree connectivity by AUROC value for each of the GCNs (COO300, HRR300, COO100 and HRR100) in all the datasets used for network annotation (CObp, GOmf, GOcc, Pfam, KEEG, PANTHER and Mapman).

### MF subnetwork was enriched in 33 terms

*PpPG21 and PpPG22 were annotated using GObp, GOmf, GOcc, Pfam, KEGG, PANTHER and MapMan. Both genes shared several annotations: ‘GOcc: extracellular region’, ‘GOcc: cell wall’, ‘GObp: metabolic process’, ‘GObp: cell wall organization’, ‘GOmf: hydrolase activity, acting on glycosyl bounds’, ‘GOmf: polygalacturonase activity’, ‘Mapman: enzyme classification. hydrolases. Glycoxylases’ and ‘Pfam: glycosyl hydrolases family 28’. PpPG22 only had two terms not shared with PpPG21, ‘GObp: fruit ripening’ and ‘GObp: carbohydrate metabolic process’*.

The *PpPG21* and *PpPG22* subnetworks were constituted by 485 and 354 genes, respectively. Even if *PpPG21* and *PpPG22* were not coexpressed, both subnetworks shared 238 genes. These genes were selected and named the melting flesh (MF) subnetwork (Figure 4; Supplementary Material 2). This MF subnetwork was annotated in GObp, GOcc, GOmf and Mapman datasets. Of the 238 genes in the MF subnetwork, 136 were annotated in *GObp*, 123 in *GOcc*, 156 in *GO*mf and 116 in Mapman (Supplementary Material 2).

**Figure 4.**
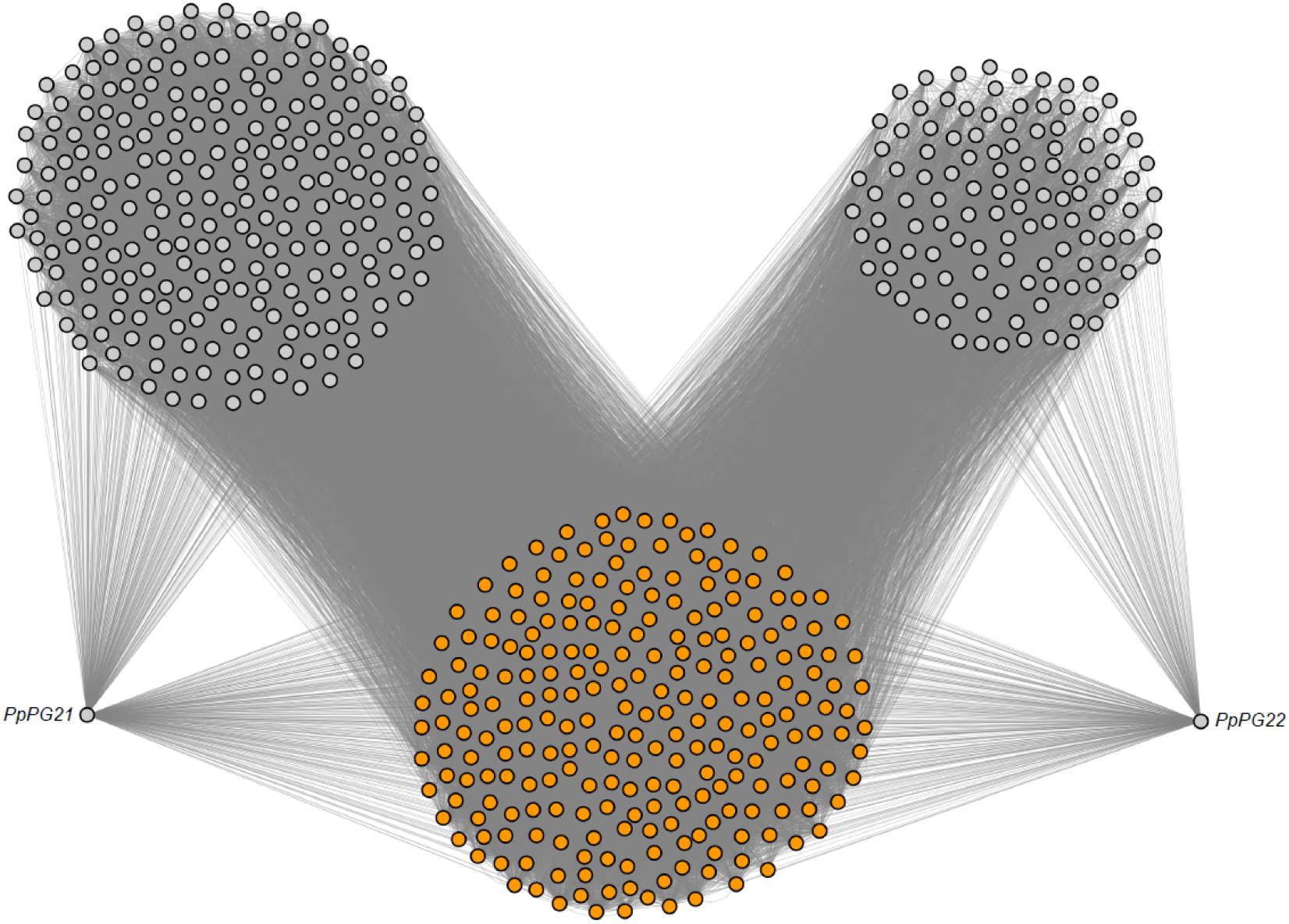
PpPG21, PpPG22 and MF subnetworks. MF subnetwork is highlighted in orange.

After MF subnetwork annotation, we performed an enrichment analysis. The MF subnetwork was enriched in 33 different terms, so 33 terms were significantly over-represented in this subnetwork. Of these 33 terms, 12 belonged to the *GOmf* dataset, nine to Mapman, eight to *GObp* and four to *GOcc* (Figure 5; Supplementary Material 2).

**Figure 5.**
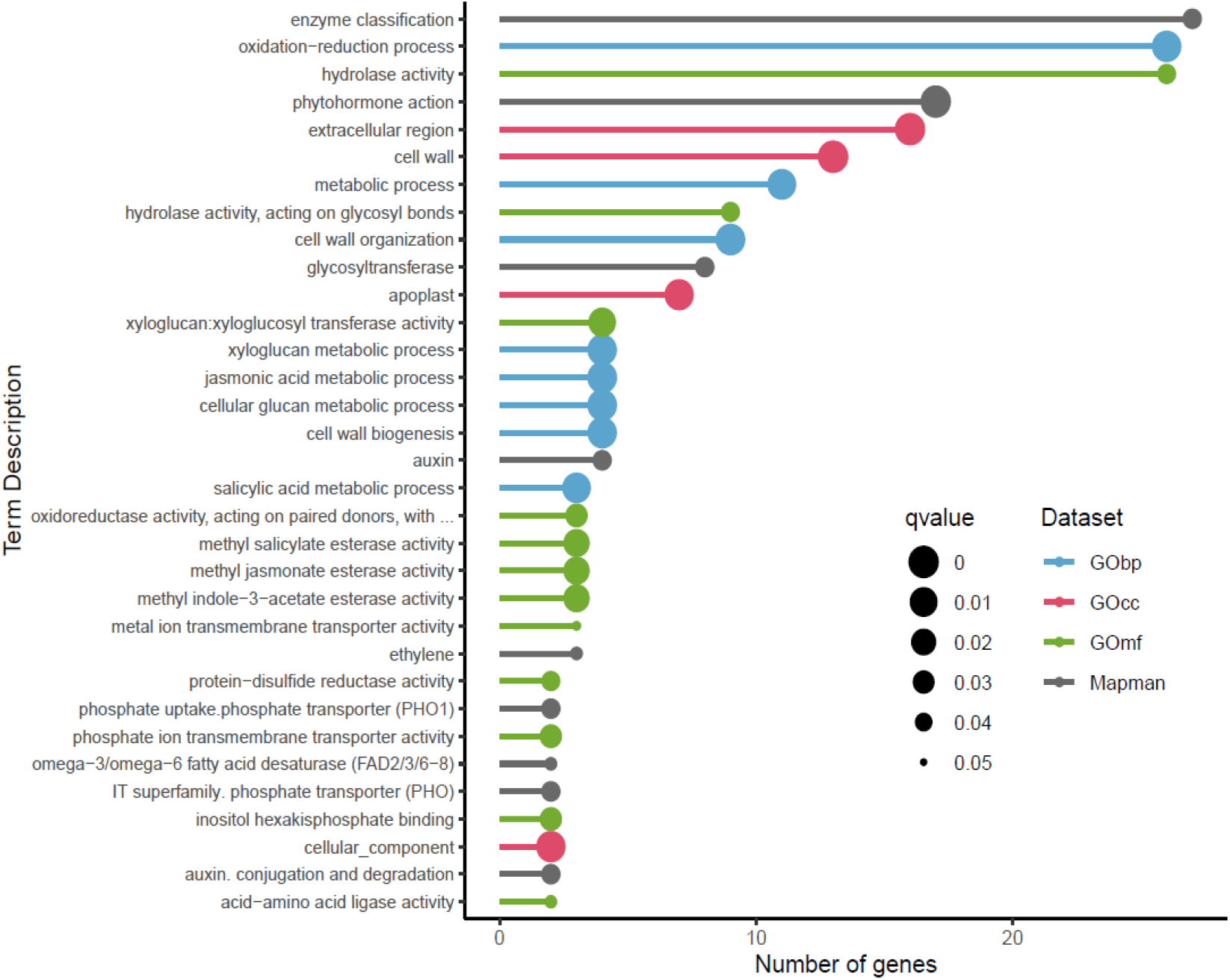
Lollipop plot of enriched terms found in the MF subnetwork. Enriched terms were sorted by the number of genes annotated by each term.

Within GOmf, up to 26 genes were annotated as *hydrolase activity* or as its child term (*direct descendant*), *hydrolase activity, acting on glycosyl bonds*. The next term was *‘xyloglucan:xyloglucosyl transferase activity’*, with four genes annotated. With three genes annotated, *we found the terms ‘methyl indole-3-acetate esterase activity’, ‘methyl salicylate esterase activity’, ‘methyl jasmonate esterase activity’, ‘oxidoreductase activity, acting on paired donors, with oxidation of a pair of donors resulting in the reduction of molecular oxygen to two molecules of water* and *metal ion transmembrane transporter activity’*. Finally, with two genes annotated, we found the terms ‘*inositol hexakisphosphate binding’*, ‘*phosphate ion transmembrane transporter activity’*, ‘*protein-disulfide reductase activity’* and ‘*acid-amino acid ligase activity’*.

Using Mapman as the annotation dataset, 27 genes were annotated as *‘enzyme classification’*. There were eight genes annotated as *‘glycosyltransferase’*, a child term of ‘*enzyme classification’*. The next term, with 17 genes annotated, was ‘*phytohormone action’*. There were four genes annotated as ‘*auxin’* or ‘*auxin*.*conjugation and degradation’* and three as ‘*ethylene’*, child terms of ‘*phytohormone action’*. With two genes annotated, we found the terms ‘*Solute transport*.*carrier-mediated transport*.*IT superfamily*.*phosphate transporter (PHO)’*, ‘*Nutrient uptake*.*phosphorus assimilation*.*phosphate uptake*.*phosphate transporter (PHO1)’* and ‘*Lipid metabolism*.*fatty acid biosynthesis*.*fatty acid desaturation*.*omega-3/omega-6 fatty acid desaturase (FAD2/3/6-8)’*.

Within GObp, there were 26 genes annotated as *‘oxidation-reduction process’*. Up to 11 genes were annotated as ‘*metabolic process’*. There were nine genes annotated as ‘*cell wall organization’* and four as ‘*cell wall biogenesis’*, child terms of *‘cell wall organization or biogenesis’*. There were four genes annotated as ‘*cellular glucan metabolic process’* and its child term, ‘*xyloglucan metabolic process’*, four as ‘*jasmonic acid metabolic process’* and three as ‘*salicylic acid metabolic process’*.

Using GOcc as the annotation dataset, 16 genes were annotated as *‘extracellular region’*, seven genes as *‘apoplast’*, child term of *‘extracellular region’*, and up to 13 genes were annotated as *‘cell wall’*.

Among the 20 negative control subnetworks created, the mean number of enriched terms was 1.35, while the melting subnetwork had 33 enriched terms (Supplementary Material 3).

## DISCUSSION

### The GCN topological characteristics are affected by the different algorithms used

To achieve the best results during gene coexpression networks (GCNs) building, two variables were tuned, aggregation method and sparsity threshold. The four GCNs obtained were evaluated, with substantial differences in the general topological characteristics of the GCNs inferred.

When considering GCN building methods, a major difference between aggregated and non-aggregated GCNs was the number of genes forming the network. Aggregated GCNs had 21,956 genes (81.7 % of *P. persica* genes), while non-aggregated GCNs only had 17,505 (65.1 % of *P. persica genes*). This difference comes from the low-expression gene filtering. In non-aggregated GCNs all the genes with less than 0.5 FPKM in 50% of the 498 RNA-Seq libraries were filtered, while in aggregated GCNs this filtering is independently performed for each of the 26 Bioproject groups. That allowed the inclusion in the GCN of genes expressed in more specific conditions and therefore involved in more specific processes. This indicates that both aggregated and non-aggregated networks were able to capture stable gene-gene relationships expressed in most of the RNA-Seq libraries used in the analysis, but only aggregated GCNs were able to detect gene-gene interactions produced in specific conditions. Condition-independent gene-gene connections could be related to basal metabolic pathways, while condition-dependent gene-gene interactions could be associated to specific metabolic pathways.

This could explain the difference in the distribution of node degree connectivity between aggregated and non-aggregated GCNs. As shown in Figure 1, aggregated GCNs had a bimodal distribution of node degree connectivity, while non-aggregated GCNs had a unimodal distribution. As mentioned previously, aggregated GCNs may be able to detect genes involved in specific and basal metabolic processes. The two modes detected in aggregated GCNs node degree connectivity distribution could be associated with these two groups of genes. The group with the lower node degree distribution could be associated with genes involved in more specific metabolic pathways, coexpressed with a lower number of genes. The group with the higher node degree distribution could be associated with genes involved in basal metabolic pathways and coexpressed with a higher number of genes. On the other hand, non-aggregated GCNs may only detect genes involved in basal metabolic pathways, having only one mode in their node degree distribution.

Another factor affecting the topology of the networks was the sparsity threshold selected. HRR300 and COO300 had a node degree connectivity higher than HRR100 and COO100. This was an expected result, since a higher number of ranked genes allows a higher number of connections between genes.

### Sparsity threshold and the number of Bioprojects determine network performance

According to the results, sparsity was a key factor affecting network performance. The average AUROC of relaxed sparsity threshold networks (HRR300 and COO300) was 0.741, while that of stringent sparsity threshold networks (HRR100 and COO100) was 0.694. Applying relaxed sparsity threshold during network building represented an increment of 6.3% in the AUROC score in comparison to stringent sparsity threshold.

The number of Bioprojects used to build the GCN was a key factor in the case of aggregated methods, indicating the minimum number of Bioprojects necessary to reach a sufficiently high AUROC score (Figure 1). In every case, aggregated methods had a lower AUROC value than non-aggregated methods using a low number of Bioprojects. By increasing this number, aggregated methods overtook non-aggregated methods, as found in other studies (Orduña et al., 2022; Orduña-Rubio et al., 2023). For future GCNs construction, increasing the number of Bioprojects could improve the performance of the GCNs.

Studying the effect of functional annotations average node degree on the AUROC value, we found major differences depending on the type of dataset used. There was a positive correlation between functional annotations average node degree and functional annotations individual AUROC in datasets based on evidence such as GObp, GOmf, GOcc, KEGG and Mapman. On the other hand, this correlation was lost with datasets based on domain identification by sequence similarity, such as PANTHER and Pfam. These results are in agreement with the GBA principle, which states that coexpressed genes share function, and not necessarily similar sequences.

### COO300 validated as a powerful tool for peach and Prunus research

In peach, fruit flesh softening has been extensively studied at fruit ripening and postharvest due to its implication in fruit shelf life. Fruit softening involves several cellular processes, such as the disassembly of the cell wall and the dissolution of the middle lamella. These modifications are the result of hydrolytic changes in the polysaccharides forming the cell wall, including celluloses, hemicelluloses (mainly xyloglucan) and pectins. Several terms found in the MF subnetwork were associated to this process, such as ‘*GOcc: cell wall’*, ‘*GObp: cell wall organization’* and ‘*GObp: cell wall biogenesis’*, ‘*GOmf: hydrolase activity’*, ‘*GOmf: hydrolase activity, actin on glycosyl bonds’*, ‘*Mapman: enzyme classification*.*EC_2 transferases*.*EC_2*.*4 glycosyltransferase’* and ‘*GOmf: xyloglucan:xyloglucosyl transferase activity’*.

Peach flesh softening is a synergistic process triggered by an extensive phytohormone signaling network. As a climacteric fruit, cross talk between ethylene and auxin occurs during peach ripening (Trainotti, Tadiello, & Casadoro, 2007). Moreover, methyl jasmonates (MeJAs) play an important role in slowing down fruit ripening by inhibiting ethylene production and fruit flesh softening (Soto, Ruiz, Ziosi, Costa, & Torrigiani, 2012; Wei, Wen, & Tang, 2017). Up to seven enriched terms were related to these phytohormones in the MF subnetwork, such as *‘Mapman: phytohormone action’, ‘Mapman: phytohormone action. Auxin’*, ‘*GObp: jasmonic acid metabolic process’*, ‘*Mapman: phytohormone action. ethylene’, ‘GOmf: methyl indole-3-esterase activity’, ‘GOmf: methyl jasmonate esterase activity’* and *‘Mapman: phytohormone action. auxin. auxin conjugation and degradation’*.

We found 25 genes in the MF subnetwork that have previously been reported as associated to ripening and softening (Supplementary Material 2). Among them, we identified several genes involved in the enzymatic machinery responsible for cell wall disassembly, such as a pectin methylesterase (*Prupe*.*7G192800*), a pectin methylesterase inhibitor (*Prupe*.*1G114500*), a pectate lyase (*Prupe*.*4G116600*), a β-galactosidase (*Prupe*.*3G050200*) and a xyloglucan endotransglycosylase hydrolase (*Prupe*.*1G255100*). Additionally, we found an expansin, a cell wall structural protein (*Prupe*.*6G075100*). Related to ethylene, we identified a 1-amino-cyclopropane-1-carboxylate synthase (*PpACS1, Prupe*.*2G176900*) and 1-amino-cyclopropane-1-carboxylate oxidase (*PpACO1, Prupe*.*3G209900*), both genes codifying the key enzymes catalyzing the final steps of the ethylene biosynthetic pathway (Tonutti et al., 1997). In fact, *PpACS1* has been previously reported as a regulator of *PpPG21* (Tatsuki et al., 2013). Another gene related to ethylene production was an ethylene receptor 2 (*PpETR2, Prupe*.*1G034300*). The implication of this gene in the ethylene transduction signal has been verified at the transcriptional level in the final stages of fruit ripening in melting flesh peaches (Wang et al., 2017). Regarding genes related to auxin biosynthesis, we found a YUCCA-like auxin-biosynthesis gene (*PpYUC11, Prupe*.*6G157500*) and an IAA amino acid synthase (*PpGH3, Prupe*.*6G226100*). Both genes have been reported to have the same expression pattern as *PpACS1* at late ripening stages in response to high auxins levels in melting flesh fruits (Pan et al., 2015).

Based on these results, we can affirm that the MF subnetwork is mainly formed by genes involved in cell wall organization and biogenesis, with expression regulated by ripening-related phytohormones such as ethylene, auxin and MeJA. Moreover, we found 25 genes previously reported as involved in softening, some taking part in key steps of these processes. These results demonstrate that the MF subnetwork is closely related to peach fruit softening and therefore to the function of *PpPG21* and *PpPG22*. Taken together, this validates COO300 as an accurate and powerful tool for peach and *Prunus* research.

### Gene coexpression networks as catalysts for Prunus research

While large-scale GCNs have been unexplored as tools in *Prunus* research until now, they are widely used in the model organism *Arabidopsis thaliana* and other crop species. Depending on the needs of the researcher, GCNs have been exploited in different ways. One of the most common is to identify different modules (also known as clusters) within the GCN through a clusterization analysis. These gene modules, which represent groups of genes highly connected between them and relatively isolated from the rest of the GCN, are particularly useful to study uncharacterized biological processes. For example, Childs et al., 2011 used this approach in rice to annotate 13,537 genes, 2,980 of which had no previous annotation.

Another approach that uses group of genes to study specific biological processes is the guide gene analysis. In this case, a list of well-characterized genes involved in a specific biological process are selected and genes coexpressing with the list of genes of interest are extracted from the network. In this way, the selected genes are used as a guide to study the transcriptional regulation of the biological process of interest. Huang et al., 2017 successfully applied this approach to study the cell wall biosynthesis in maize. Pathway-centered network analysis has also been helpful in the identification of members or regulators of secondary metabolic pathways (Orduña-Rubio et al., 2023).

GCNs can also be used to study specific gene families, being particularly useful for studying transcription factor families. For instance, Wong et al., 2016 developed a MYB-centered GCN to study the potential processes being regulated by this family in grapevine.

Finally, GCNs can be used to infer the function of a gene of interest. This is a situation of special interest in peach and *Prunus* research, where most trait-loci analyses lead to a list of candidate genes associated with the trait under study. With poor or no functional information, identifying the responsible gene from this list of candidates can be almost impossible. Even when a high-confidence candidate gene is identified, the lack of an efficient genetic transformation system is still one of the main limitations for functional, mutant, or transgenic based validation. Having a tool such as the GCN presented in this study, with which obtaining useful information about the biological processes in which a gene is involved, may be of critical importance.

## CONCLUSIONS

In this study, we performed the widest overview of transcriptomic analysis carried out to date in peach or other *Prunus species*. The GCN inference methods used, aggregated or non-aggregated, affected the topological characteristics and performance of the GCNs created. Using two well-characterized genes in peach, *PpPG21* and *PpPG22*, we were able to validate the network with the best performance, COO300. The GCN tool presented in this study will help *Prunus* researchers overcome the intrinsic limitations of working with crop tree species, prioritize research lines and outline new ones. COO300, named as PeachGCN v1.0, and the scripts necessary to run a function prediction analysis using it, are available at https://github.com/felipecobos/PeachGCN.

## BIBLIOGRAPHY

Amrine, K. C. H., Blanco-Ulate, B., & Cantu, D. (2015). Discovery of Core Biotic Stress Responsive Genes in Arabidopsis by Weighted Gene Co-Expression Network Analysis. PLOS ONE, 10(3), e0118731. https://doi.org/10.1371/JOURNAL.PONE.0118731

Aranzana, M. J., Decroocq, V., Dirlewanger, E., Eduardo, I., Gao, Z. S., Gasic, K., Iezzoni, A., Jung, S., Peace, C., Prieto, H., Tao, R., Verde, I., Abbott, A. G., & Arús, P. (2019). Prunus genetics and applications after de novo genome sequencing: achievements and prospects. In Horticulture Research (Vol. 6, Issue 1, p. 58). Nature Publishing Group. https://doi.org/10.1038/s41438-019-0140-8

Ashburner, M., Ball, C. A., Blake, J. A., Botstein, D., Butler, H., Cherry, J. M., Davis, A. P., Dolinski, K., Dwight, S. S., Eppig, J. T., Harris, M. A., Hill, D. P., Issel-Tarver, L., Kasarskis, A., Lewis, S., Matese, J. C., Richardson, J. E., Ringwald, M., Rubin, G. M., & Sherlock, G. (2000). Gene Ontology: tool for the unification of biology. Nature Genetics 2000 25:1, 25(1), 25–29. https://doi.org/10.1038/75556

Ballouz, S., Weber, M., Pavlidis, P., & Gillis, J. (2017). EGAD: Ultra-fast functional analysis of gene networks. Bioinformatics, 33(4), 612–614. https://doi.org/10.1093/bioinformatics/btw695

Carbon, S., Douglass, E., Good, B. M., Unni, D. R., Harris, N. L., Mungall, C. J., Basu, S., Chisholm, R. L., Dodson, R. J., Hartline, E., Fey, P., Thomas, P. D., Albou, L.-P., Ebert, D., Kesling, M. J., Mi, H., Muruganujan, A., Huang, X., Mushayahama, T., & LaBonte, S. A. (2021). The Gene Ontology resource: enriching a GOld mine [Article]. Nucleic Acids Research., 49(D1), D325–D334. https://doi.org/10.1093/nar/gkaa1113

Cheng, C., Liu, J., Wang, X., Wang, Y., Yuan, Y., & Yang, S. (2022). PpERF/ABR1 functions as an activator to regulate PpPG expression resulting in fruit softening during storage in peach (Prunus persica). Postharvest Biology and Technology, 189, 111919. https://doi.org/10.1016/J.POSTHARVBIO.2022.111919

Childs, K. L., Davidson, R. M., & Buell, C. R. (2011). Gene Coexpression Network Analysis as a Source of Functional Annotation for Rice Genes. PLoS ONE, 6(7), e22196. https://doi.org/10.1371/journal.pone.0022196

Danecek, P., Bonfield, J. K., Liddle, J., Marshall, J., Ohan, V., Pollard, M. O., Whitwham, A., Keane, T., McCarthy, S. A., Davies, R. M., & Li, H. (2021). Twelve years of SAMtools and BCFtools. GigaScience, 10(2), 1–4. https://doi.org/10.1093/GIGASCIENCE/GIAB008

Dardick, C., Callahan, A., Horn, R., Ruiz, K. B., Zhebentyayeva, T., Hollender, C., Whitaker, M., Abbott, A., & Scorza, R. (2013). PpeTAC1 promotes the horizontal growth of branches in peach trees and is a member of a functionally conserved gene family found in diverse plants species. The Plant Journal, 75(4), 618–630. https://doi.org/10.1111/TPJ.12234

Durinck, S., Spellman, P. T., Birney, E., & Huber, W. (2009). Mapping identifiers for the integration of genomic datasets with the R/Bioconductor package biomaRt. Nature Protocols 2009 4:8, 4(8), 1184–1191. https://doi.org/10.1038/nprot.2009.97

Ficklin, S. P., Luo, F., & Feltus, F. A. (2010). The Association of Multiple Interacting Genes with Specific Phenotypes in Rice Using Gene Coexpression Networks. Plant Physiology, 154(1), 13–24. https://doi.org/10.1104/PP.110.159459

Furuya, T., Saito, M., Uchimura, H., Satake, A., Nosaki, S., Miyakawa, T., Shimadzu, S., Yamori, W., Tanokura, M., Fukuda, H., & Kondo, Y. (2021). Gene co-expression network analysis identifies BEH3 as a stabilizer of secondary vascular development in Arabidopsis. The Plant Cell. https://doi.org/10.1093/PLCELL/KOAB151

García-Gómez, B. E., Ruiz, D., Salazar, J. A., Rubio, M., Martínez-García, P. J., & Martínez-Gómez, P. (2020). Analysis of Metabolites and Gene Expression Changes Relative to Apricot (Prunus armeniaca L.) Fruit Quality During Development and Ripening. Frontiers in Plant Science, 0, 1269. https://doi.org/10.3389/FPLS.2020.01269

Gu, C., Wang, L., Wang, W., Zhou, H., Ma, B., Zheng, H., Fang, T., Ogutu, C., Vimolmangkang, S., & Han, Y. (2016). Copy number variation of a gene cluster encoding endopolygalacturonase mediates flesh texture and stone adhesion in peach. Journal of Experimental Botany, 67(6), 1993–2005. https://doi.org/10.1093/JXB/ERW021

Guseman, J. M., Webb, K., Srinivasan, C., & Dardick, C. (2017). DRO1 influences root system architecture in Arabidopsis and Prunus species. The Plant Journal, 89(6), 1093–1105. https://doi.org/10.1111/TPJ.13470

Huang, J., Vendramin, S., Shi, L., & McGinnis, K. M. (2017). Construction and optimization of a large gene coexpression network in maize using RNA-seq data. Plant Physiology, 175(1), 568–583. https://doi.org/10.1104/pp.17.00825

Jiang, L., Kang, R., Feng, L., Yu, Z., & Luo, H. (2020). iTRAQ-based quantitative proteomic analysis of peach fruit (Prunus persica L.) at different ripening and postharvest storage stages. Postharvest Biology and Technology, 164, 111137. https://doi.org/10.1016/J.POSTHARVBIO.2020.111137

Jiang, X., Liu, K., Peng, H., Fang, J., Zhang, A., Han, Y., & Zhang, X. (2023). Comparative network analysis reveals the dynamics of organic acid diversity during fruit ripening in peach (Prunus persica L. Batsch). BMC Plant Biology, 23(1), 1–14. https://doi.org/10.1186/S12870-023-04037-W/TABLES/1

Jung, S., Lee, T., Cheng, C.-H., Buble, K., Zheng, P., Yu, J., Humann, J., Ficklin, S. P., Gasic, K., Scott, K., Frank, M., Ru, S., Hough, H., Evans, K., Peace, C., Olmstead, M., DeVetter, L. W., McFerson, J., Coe, M., … Main, D. (2019). 15 years of GDR: New data and functionality in the Genome Database for Rosaceae. Nucleic Acids Research, 47(D1), D1137–D1145. https://doi.org/10.1093/nar/gky1000

Kanehisa, M., & Goto, S. (2000). KEGG: Kyoto Encyclopedia of Genes and Genomes. Nucleic Acids Research, 28(1), 27–30. https://doi.org/10.1093/NAR/28.1.27

Kim, D., Langmead, B., & Salzberg, S. L. (2015). HISAT: a fast spliced aligner with low memory requirements. Nature Methods 2015 12:4, 12(4), 357–360. https://doi.org/10.1038/nmeth.3317

Leinonen, R., Sugawara, H., & Shumway, M. (2011). The sequence read archive. Nucleic Acids Research, 39(SUPPL. 1), D19–D21. https://doi.org/10.1093/nar/gkq1019

Li, H., Handsaker, B., Wysoker, A., Fennell, T., Ruan, J., Homer, N., Marth, G., Abecasis, G., & Durbin, R. (2009). The Sequence Alignment/Map format and SAMtools. Bioinformatics, 25(16), 2078–2079. https://doi.org/10.1093/bioinformatics/btp352

Liao, Y., Smyth, G. K., & Shi, W. (2014). featureCounts: an efficient general purpose program for assigning sequence reads to genomic features. Bioinformatics, 30(7), 923–930. https://doi.org/10.1093/BIOINFORMATICS/BTT656

Limera, C., Sabbadini, S., Sweet, J. B., & Mezzetti, B. (2017). New Biotechnological Tools for the Genetic Improvement of Major Woody Fruit Species. Frontiers in Plant Science, 0, 1418. https://doi.org/10.3389/FPLS.2017.01418

Liu, W., Lin, L., Zhang, Z., Liu, S., Gao, K., Lv, Y., Tao, H., & He, H. (2019). Gene co-expression network analysis identifies trait-related modules in Arabidopsis thaliana. Planta 2019 249:5, 249(5), 1487–1501. https://doi.org/10.1007/S00425-019-03102-9

Lv, L., Zhang, W., Sun, L., Zhao, A., Zhang, Y., Wang, L., Liu, Y., Li, Z., Li, H., & Chen, X. (2020). Gene co-expression network analysis to identify critical modules and candidate genes of drought-resistance in wheat. PLOS ONE, 15(8), e0236186. https://doi.org/10.1371/JOURNAL.PONE.0236186

Ma, S., Ding, Z., & Li, P. (2017). Maize network analysis revealed gene modules involved in development, nutrients utilization, metabolism, and stress response. BMC Plant Biology, 17(1), 1–17. https://doi.org/10.1186/s12870-017-1077-4

Mao, L., Van Hemert, J. L., Dash, S., & Dickerson, J. A. (2009). Arabidopsis gene co-expression network and its functional modules. BMC Bioinformatics 2009 10:1, 10(1), 1–24. https://doi.org/10.1186/1471-2105-10-346

Mi, H., Ebert, D., Muruganujan, A., Mills, C., Albou, L.-P., Mushayamaha, T., & Thomas, P. D. (2021). PANTHER version 16: a revised family classification, tree-based classification tool, enhancer regions and extensive API. Nucleic Acids Research, 49(D1), D394–D403. https://doi.org/10.1093/NAR/GKAA1106

Mistry, J., Chuguransky, S., Williams, L., Qureshi, M., Salazar, G. A., Sonnhammer, E. L. L., Tosatto, S. C. E., Paladin, L., Raj, S., Richardson, L. J., Finn, R. D., & Bateman, A. (2021). Pfam: The protein families database in 2021. Nucleic Acids Research, 49(D1), D412–D419. https://doi.org/10.1093/NAR/GKAA913

Nakano, R., Kawai, T., Fukamatsu, Y., Akita, K., Watanabe, S., Asano, T., Takata, D., Sato, M., Fukuda, F., & Ushijima, K. (2020). Postharvest Properties of Ultra-Late Maturing Peach Cultivars and Their Attributions to Melting Flesh (M) Locus: Re-evaluation of M Locus in Association With Flesh Texture. Frontiers in Plant Science, 11, 1817. https://doi.org/10.3389/FPLS.2020.554158/BIBTEX

Oliver, S. (2000). Guilt-by-association goes global. Nature 2000 403:6770, 403(6770), 601–602. https://doi.org/10.1038/35001165

Orduña, L., Li, M., Navarro-Payá, D., Zhang, C., Santiago, A., Romero, P., Ramšak, Ž., Magon, G., Höll, J., Merz, P., Gruden, K., Vannozzi, A., Cantu, D., Bogs, J., Wong, D. C. J., Huang, S. shan C., & Matus, J. T. (2022). Direct regulation of shikimate, early phenylpropanoid, and stilbenoid pathways by Subgroup 2 R2R3-MYBs in grapevine. Plant Journal, 110(2), 529–547. https://doi.org/10.1111/TPJ.15686

Orduña-Rubio, L., Santiago, A., Navarro-Payá, D., Zhang, C., Wong, D. C. J., & Matus, J. T. (2023). Aggregated gene co-expression networks for predicting transcription factor regulatory landscapes in a non-model plant species. BioRxiv. https://doi.org/https://doi.org/10.1101/2023.04.24.538042

Pan, L., Zeng, W., Niu, L., Lu, Z., Liu, H., Cui, G., Zhu, Y., Chu, J., Li, W., Fang, W., Cai, Z., Li, G., & Wang, Z. (2015). PpYUC11, a strong candidate gene for the stony hard phenotype in peach (Prunus persica L. Batsch), participates in IAA biosynthesis during fruit ripening. Journal of Experimental Botany, 66(22), 7031–7044. https://doi.org/10.1093/JXB/ERV400

Qian, M., Xu, Z., Zhang, Z., Li, Q., Yan, X., Liu, H., Han, M., Li, F., Zheng, J., Zhang, D., & Zhao, C. (2021). The downregulation of PpPG21 and PpPG22 influences peach fruit texture and softening. Planta 2021 254:2, 254(2), 1–12. https://doi.org/10.1007/S00425-021-03673-6

Ricci, A., Sabbadini, S., Prieto, H., Padilla, I. M., Dardick, C., Li, Z., Scorza, R., Limera, C., Mezzetti, B., Perez-Jimenez, M., Burgos, L., & Petri, C. (2020). Genetic Transformation in Peach (Prunus persica L.): Challenges and Ways Forward. Plants 2020, Vol. 9, Page 971, 9(8), 971. https://doi.org/10.3390/PLANTS9080971

Sayers, E. W., Bolton, E. E., Brister, J. R., Canese, K., Chan, J., Comeau, D. C., Connor, R., Funk, K., Kelly, C., Kim, S., Madej, T., Marchler-Bauer, A., Lanczycki, C., Lathrop, S., Lu, Z., Thibaud-Nissen, F., Murphy, T., Phan, L., Skripchenko, Y., … Sherry, S. T. (2022a). Database resources of the national center for biotechnology information. Nucleic Acids Research, 50(D1), D20–D26. https://doi.org/10.1093/NAR/GKAB1112

Sayers, E. W., Bolton, E. E., Brister, J. R., Canese, K., Chan, J., Comeau, D. C., Connor, R., Funk, K., Kelly, C., Kim, S., Madej, T., Marchler-Bauer, A., Lanczycki, C., Lathrop, S., Lu, Z., Thibaud-Nissen, F., Murphy, T., Phan, L., Skripchenko, Y., … Sherry, S. T. (2022b). Database resources of the national center for biotechnology information. Nucleic Acids Research, 50(D1), D20–D26. https://doi.org/10.1093/NAR/GKAB1112

Schaefer, R. J., Michno, J. M., & Myers, C. L. (2017). Unraveling gene function in agricultural species using gene co-expression networks. Biochimica et Biophysica Acta (BBA) - Gene Regulatory Mechanisms, 1860(1), 53–63. https://doi.org/10.1016/J.BBAGRM.2016.07.016

Soto, A., Ruiz, K. B., Ziosi, V., Costa, G., & Torrigiani, P. (2012). Ethylene and auxin biosynthesis and signaling are impaired by methyl jasmonate leading to a transient slowing down of ripening in peach fruit. Journal of Plant Physiology, 169(18), 1858–1865. https://doi.org/10.1016/J.JPLPH.2012.07.007

Thimm, O., Bläsing, O., Gibon, Y., Nagel, A., Meyer, S., Krüger, P., Selbig, J., Müller, L. A., Rhee, S. Y., & Stitt, M. (2004). mapman: a user-driven tool to display genomics data sets onto diagrams of metabolic pathways and other biological processes. The Plant Journal, 37(6), 914–939. https://doi.org/10.1111/J.1365-313X.2004.02016.X

Verde, I., Abbott, A. G., Scalabrin, S., Jung, S., Shu, S., Marroni, F., Zhebentyayeva, T., Dettori, M. T., Grimwood, J., Cattonaro, F., Zuccolo, A., Rossini, L., Jenkins, J., Vendramin, E., Meisel, L. A., Decroocq, V., Sosinski, B., Prochnik, S., Mitros, T., … Rokhsar, D. S. (2013). The high-quality draft genome of peach (Prunus persica) identifies unique patterns of genetic diversity, domestication and genome evolution. Nature Genetics, 45(5), 487–494. https://doi.org/10.1038/ng.2586

Verde, I., Jenkins, J., Dondini, L., Micali, S., Pagliarani, G., Vendramin, E., Paris, R., Aramini, V., Gazza, L., Rossini, L., Bassi, D., Troggio, M., Shu, S., Grimwood, J., Tartarini, S., Dettori, M. T., & Schmutz, J. (2017). The Peach v2.0 release: High-resolution linkage mapping and deep resequencing improve chromosome-scale assembly and contiguity. BMC Genomics, 18(1). https://doi.org/10.1186/s12864-017-3606-9

Wang, Q., Cao, K., Li, Y., Wu, J., Fan, J., Ding, T., Khan, I. A., & Wang, L. (2023). Identification of co-expressed networks and key genes associated with organic acid in peach fruit. Scientia Horticulturae, 307, 111496. https://doi.org/10.1016/j.scienta.2022.111496

Wang, X., Ding, Y., Wang, Y., Pan, L., Niu, L., Lu, Z., Cui, G., Zeng, W., & Wang, Z. (2017). Genes involved in ethylene signal transduction in peach (Prunus persica) and their expression profiles during fruit maturation. Scientia Horticulturae, 224, 306–316. https://doi.org/10.1016/J.SCIENTA.2017.06.035

Wang, Z., Gerstein, M., & Snyder, M. (2009). RNA-Seq: a revolutionary tool for transcriptomics. Nature Reviews Genetics 2008 10:1, 10(1), 57–63. https://doi.org/10.1038/nrg2484

Wong, D. C. J. (2020). Network aggregation improves gene function prediction of grapevine gene co-expression networks. Plant Molecular Biology, 103(4–5), 425–441. https://doi.org/10.1007/s11103-020-01001-2

Wong, D. C. J., Schlechter, R., Vannozzi, A., Höll, J., Hmmam, I., Bogs, J., Tornielli, G. B., Castellarin, S. D., & Matus, J. T. (2016). A systems-oriented analysis of the grapevine R2R3-MYB transcription factor family uncovers new insights into the regulation of stilbene accumulation. DNA Research, 23(5), 451–466. https://doi.org/10.1093/dnares/dsw028

Wu, X., Du, A., Zhang, S., Wang, W., Liang, J., Peng, F., & Xiao, Y. (2021). Regulation of growth in peach roots by exogenous hydrogen sulfide based on RNA-Seq. Plant Physiology and Biochemistry, 159, 179–192. https://doi.org/10.1016/J.PLAPHY.2020.12.018

Xi, W., Feng, J., Liu, Y., Zhang, S., & Zhao, G. (2019). The R2R3-MYB transcription factor PaMYB10 is involved in anthocyanin biosynthesis in apricots and determines red blushed skin. BMC Plant Biology, 19(1). https://doi.org/10.1186/s12870-019-1898-4

Zhang, Q., Feng, C., Li, W., Qu, Z., Zeng, M., & Xi, W. (2019). Transcriptional regulatory networks controlling taste and aroma quality of apricot (Prunus armeniaca L.) fruit during ripening. BMC Genomics, 20(1), 45. https://doi.org/10.1186/s12864-019-5424-8

Zhu, Y., Zeng, W., Wang, X., Pan, L., Niu, L., Lu, Z., Cui, G., & Wang, Z. (2017). Characterization and Transcript Profiling of PME and PMEI Gene Families during Peach Fruit Maturation. Journal of the American Society for Horticultural Science, 142(4), 246–259. https://doi.org/10.21273/JASHS04039-17

