## Supplementary figures and images for "First large-scale peach gene coexpression network: A new tool for predicting gene function"

### Supplementary Figure 1

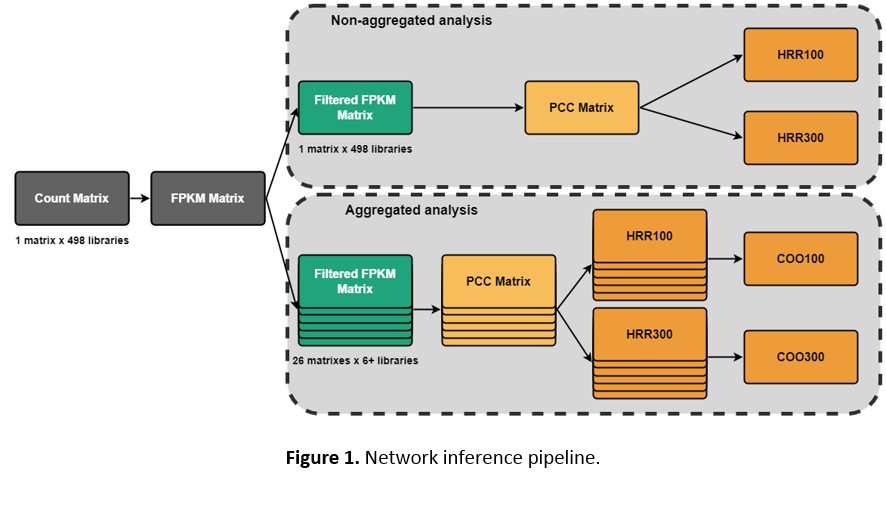
